# Reconstructor: A COBRApy compatible tool for automated genome-scale metabolic network reconstruction with parsimonious flux-based gap-filling

**DOI:** 10.1101/2022.09.17.508371

**Authors:** Matthew L Jenior, Emma M Glass, Jason A Papin

## Abstract

**Summary:** Genome-scale metabolic network reconstructions (GENREs) are valuable for understanding cellular metabolism *in silico*. Several tools exist for automatic GENRE generation. However, these tools frequently (1) do not readily integrate with some of the widely-used suites of packaged methods available for network analysis, (2) lack effective network curation tools, and (3) are not sufficiently user-friendly. Here, we present Reconstructor, a user-friendly COBRApy compatible tool with ModelSEED namespace compatibility and a pFBA-based gap-filling technique. We demonstrate how Reconstructor readily generates high-quality GENRES that are useful for further biological discovery.

**Availability and Implementation:** The Reconstructor package is freely available for download via pip in the command line (pip install reconstructor). Usage instructions and benchmarking data are available at http://github.com/emmamglass/reconstructor.

**Contact:** Jason Papin

## Introduction

Genome-scale metabolic network reconstructions (GENREs) are valuable tools for understanding the link between the genotype and phenotype of an organism. GENREs enable greater understanding of the effects of genetic and environmental perturbation on cellular function and can help to identify novel drug targets, among many other applications (Haggart *et al*., 2011; Gu *et al*., 2019; Kim *et al*., 2012).

The synthesis of GENREs can be an incredibly laborious and complex process, requiring the integration of data from multiple sources (Thiele and Palsson, 2010). The creation of a GENRE begins with the annotated genome sequence to predict reactions to include in the draft GENRE, and then further model curation steps are performed to gap-fill missing reactions. While GENREs can be generated and curated manually, methods for the automated creation of GENREs have emerged (Mendoza *et al*., 2019).

Several platforms exist for automated GENRE creation, including ModelSEED (Seaver *et al*., 2021) and CarveMe (Machado *et al*., 2018), among others (Chevallier *et al*., 2018; Dias *et al*., 2015; Olivier, 2018; Karp *et al*.; Wang *et al*., 2018) (Figure 1B). However, additional compatibility modules are necessary to use CarveMe- and ModelSEED-created GENREs with the COBRApy analysis toolbox (Moretti *et al*., 2016; Mundy *et al*., 2017), due to the web-based nature of ModelSEED and the use of the BiGG reaction database by CarveMe. Additionally, conventional automated GENRE creators use gene-protein-reaction (GPR) associations for reaction scoring and subsequent gap-filling, which could lead to the inclusion of reactions in the final model that are unnecessary. GPR associations cannot be solely relied upon for gap-filling because their formulation relies on sufficient genome annotation data and experimentally obtained information, which can be lacking for certain organisms (Mendoza *et al*., 2019).

**Figure 1.**
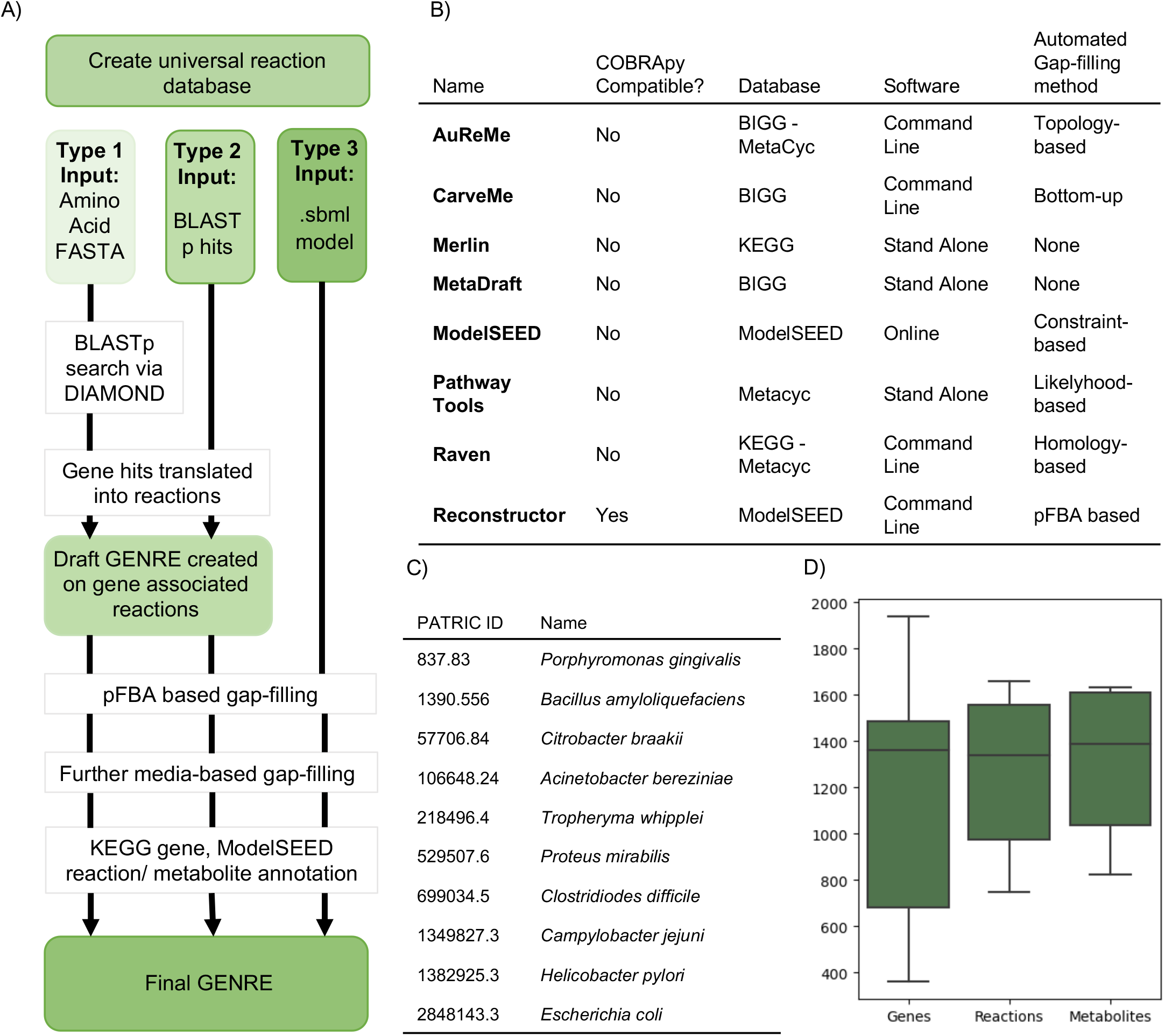
Reconstructor overview. A) Flowchart detailing the functionality of the Reconstructor tool. B) Comparison of other widely used GENRE construction including Reconstructor, adapted from (Mendoza *et al*., 2019). C) GENREs were created via Reconstructor for each of the 10 bacterial species listed, genome sequences were downloaded from the Pathosystems Resource Integration Center (PATRIC) (Davis *et al*., 2020), PATRIC IDs for each species are listed. D) Boxplots of the number of genes, reactions, and metabolites in the 10 GENREs

Here, we introduce Reconstructor, an automated GENRE creation tool that creates COBRApy-compatible reconstructions in the ModelSEED namespace. Additionally, we include a gap-filling technique based on parsimonious flux balance analysis (pFBA), a more biologically tractable gap-filing technique than other techniques based exclusively on GPR mapping.

## Results

### Universal reaction database construction

A universal database of metabolic reactions was created based on all available reactions and metabolites in the ModelSEED database. Reconstructor GENREs are in the ModelSEED namespace, but are also directly compatible with COBRApy, without needing the use of additional compatibility modules. All ModelSEED reactions and metabolites were added to the universal database via reaction and metabolite dictionaries, missing exchange reactions were identified and corrected, and biomass function was updated. It is important to note that the gram-positive and gram-negative biomass functions were differentially defined. The universal reaction database was further curated to remove mass-imbalanced reactions and reactions with no reactants. Intracellular sink reactions were added. The result was a universal database that contains a reaction collection from which the genome-informed model can select reactions for gap-filling. The user can also curate their own universal database to use with Reconstructor by altering the ModelSEED reaction and metabolite dictionaries. The ability to curate readily this provided database or to make use of any other user-provided universal database in the same name-space is a key feature of Reconstructor.

### Input data formats and draft GENRE scaffold extraction

Reconstructor automates the build of a GENRE from three different types of user defined input. Type 1 requires inputs of an annotated genome sequence in the form of an amino acid FASTA file and the gram status of the bacterial species. Type 2 requires an input of BLASTp hits and the gram status of the bacterial species, bypassing the BLASTp search step. Type 3 requires an existing GENRE in sbml format, and further gap-filling is performed based on pFBA gap-filling (as described further below). Additionally, the user can define their own media conditions for a given GENRE by providing metabolite names present in their defined media condition.

The GENRE creation process is described below from the starting point of a Type 1 input. The amino acid FASTA is aligned to the KEGG database by performing a BLASTp search with the DIAMOND sequence aligner tool (Buchfink *et al*., 2014). Then, the gene hits are processed and translated into reactions. These reactions and associated gene names are used to create a draft GENRE based solely on gene associated reactions. Additionally, reactions are added to the draft GENRE based on defined media conditions.

### pFBA-based approach to gap-filling draft GENREs

Several gap-filling methods exist (Pan and Reed, 2018), many of which use parsimony as a guiding principle in which a minimum number of reactions are added to satisfy criteria like growth in defined media (Prigent *et al*., 2017; King *et al*., 2018; Zimmermann *et al*., 2021). In Reconstructor, the draft GENRE is gap-filled by adding reactions to the draft GENRE from the universal reaction database through a pFBA approach. The pFBA gap-filler first modifies the universal reaction database by removing any reactions that are present already in the gene associated reaction draft GENRE. Then, draft GENRE reactions are added to the universal reaction database. The optimal objective flux for the draft GENRE is calculated and used to constrain further optimization to a fraction of this level of flux. Each reaction is assigned a weight in the new objective function with non-gene associated, universal reactions at a maximum weight of 1. The sum of the reaction and linear coefficient combinations for each forward and reverse reaction pair is used as the new objective function. This linear coefficient assignment forces flux through gene-associated reactions and minimizes the inclusion of non-gene associated reactions. New reactions are added to the draft GENRE if the absolute value of the solution flux is greater than 1×10^−6^.

### Secondary Gap-filling, component annotation, and GENRE output

Further gap-filling is performed based on defined media conditions. The GENRE is then annotated with KEGG genes, ModelSEED metabolites/reactions, and reactions implicated in biomass are defined. Finally, exchange reactions are corrected, and basic model checks are completed to report the number of genes, reactions, and metabolites in the draft and final GENREs, how many reactions were gap-filled, and the final objective flux. Finally, the model is saved to sbml format, the current community standard (Hucka *et al*., 2003).

### COBRApy compatibility

Current widely-used GENRE creation tools, ModelSEED and CarveMe, both require additional modules to be used in conjunction with COBRApy (Moretti *et al*., 2016; Mundy *et al*., 2017). GENREs created with Reconstructor are directly compatible with COBRApy; they can be directly imported into python after creation and do not require additional compatibility modules to take advantage of the powerful COBRApy analysis toolbox. Reconstructor’s direct COBRApy compatibility allows users to streamline automated GENRE analysis pipelines, potentially accelerating GENRE-based discovery and hypothesis generation.

### Reconstructor generates high quality reconstructions

As a demonstration of the utility of Reconstructor, 10 GENREs representing unique bacterial strains were created by Reconstructor for analysis and benchmarking through the metabolic model testing suite (MEMOTE). MEMOTE scores for each of the 10 reconstructions (Figure 1C) are available at http://github.com/emmamglass/reconstructor. Each of the 10 generated GENREs were imported into COBRApy and the number of genes, metabolites, and reactions were determined (Figure 1D). The number of genes, reactions, and metabolites present within Reconstructor-generated models is consistent with GENREs created with other tools (Machado *et al*., 2018).

## Conclusion

Reconstructor automatically creates and curates COBRApy-compatible, genome-scale metabolic network reconstructions in the ModelSEED namespace and uses a pFBA based gap-filling technique (Figure 1A) that is more efficient and consistent with parsimony principles in metabolic modeling than conventional gap-filling techniques (Jenior *et al*., 2020). Direct COBRApy compatibility enables the user to import GENREs directly into python for further downstream analysis via the robust COBRApy toolbox. Reconstructor generates high-quality GENREs as evidenced through MEMOTE benchmarking, and Reconstructor GENREs are characteristically similar to those created by other tools.

### Funding Information

This work was supported by the National Science Foundation (NSF) [1842490]; National Institutes of Health [grant numbers T32 GM 145443-1, R01-AI154242, R01-AAT010253]; and the TransUniversity Microbiome Initiative.

